# A BBX-HY5 photomorphogenic module induces anthocyanin synthesis in purple tomato fruits

**DOI:** 10.64898/2025.12.03.692022

**Authors:** Jacopo Menconi, Noemi La Monaca, Pierdomenico Perata, Silvia Gonzali

## Abstract

Over the last decades, the search for novel nutraceutical foods led to the development of several anthocyanin enriched crops, with the purple tomato being one of the most prominent examples. In non-GM purple tomato, anthocyanin biosynthesis in fruits is conferred by the introgression of specific alleles of *Anthocyanin fruit (Aft)* and *atroviolacea* (*atv*) genes, which allow the pigmentation of the epicarp under optimal environmental conditions. Light induces anthocyanin synthesis through ELONGATED HYPOCOTYL 5 (HY5), whose levels are in turn controlled by CONSTITUTIVE PHOTOMORPHOGENIC 1 (COP1)-targeted destabilization. In recent years, light-related HY5-independent factors have been object of research. In this work, we have identified two novel transcription factors, belonging to the BBX protein family, modulating anthocyanin synthesis in purple tomato fruits at different stages of the pathway, including transcriptional and post-translational levels, and showing both HY5-dependent and independent functions. Our results contribute to expanding the knowledge on the light-induced regulation of anthocyanin synthesis in tomato and may serve for the development of novel anthocyanin enriched lines able to overcome the threat posed by non-optimal light conditions.

## Introduction

Over the last decades novel nutraceutical foods, including the recently marketed purple tomatoes (*Solanum lycopersicum*), have attracted a great research interest for their positive impact on human health, putting anthocyanins and other phytochemicals they contain at the focus of attention (Glover and Martin 2012; Cappellini et al. 2021).

The basic transcriptional regulation of the anthocyanin biosynthesis in plants was characterized roughly two decades ago, but the last years have been particularly enthralling and productive in demonstrating how the complete regulation of the process is much more complex than believed. The anthocyanin biosynthesis is conserved in higher plants and the structural genes involved in the pathway are traditionally divided in two groups: the early biosynthetic genes (EBGs), coding for the enzymes acting in the early reactions which produce the intermediate molecules common to different classes of flavonoids, and the late biosynthetic genes (LBGs), encoding the enzymes catalysing the final specific steps of the anthocyanin synthesis (Cappellini et al. 2021) (Supplementary Fig. S1). Several transcription factors (TFs) control the transcription of these structural genes, whose expression is in turn regulated by environmental or developmental cues. Single R2R3-MYB TFs mainly control the expression of the EBGs, whereas R2R3-MYB, basic helix-loop-helix (bHLH), and WD-repeat (WDR) factors, interacting to form the MYB-bHLH-WDR (MBW) complex, represent the principal activators of the LBGs (Xu et al. 2015; Cappellini et al. 2021) (Supplementary Fig. S1).

The regulation of anthocyanin synthesis in non-GM purple tomato fruits is under control of environmental signals and has been subject of extensive research (Mes et al. 2008; Adato et al. 2009; Povero et al. 2011). Several studies demonstrated the role of the *Anthocyanin fruit (Aft)* and *atroviolacea* (*atv*) genetic loci, introgressed from wild Solanum species (Mes et al. 2008), in the activation of the anthocyanin pathway. The *Aft* locus contains a functional allele of *AN2like,* the gene coding for the R2R3-MYB TF which is produced in the fruit epicarp upon high light exposure, taking part to MBW complexes containing the bHLH proteins JAF13 or AN1, and the WDR factor AN11 (Kiferle et al. 2015; Qiu et al. 2016; Gao et al. 2018). *AN2like* undergoes splicing mutation in common tomato varieties preventing anthocyanin synthesis in their fruits (Colanero et al. 2019; Sun et al. 2020). The *atv* locus contains a mutant and non-functional allele of *MYB*-*ATV,* a gene coding for a R3-MYB factor which normally represses the anthocyanin synthesis with a feedback mechanism (Colanero et al. 2018; Yan et al. 2020). The *Aft* and *atv* loci synergistically interact in the *Aft*/*Aft* x *atv*/*atv* (*Aft*/*atv*) tomato lines resulting in a strong activation of the synthesis of anthocyanins which can result in almost complete pigmentation of the fruit epicarp under optimal environmental conditions (Povero et al. 2011)

Light represents a major factor inducing the anthocyanin synthesis in these fruits (Zoratti et al. 2014). Ambient temperature may further affect biosynthetic or degradation processes through upregulation or downregulation of TFs involved in the anthocyanin metabolic pathways (Kiferle et al. 2015; Qiu et al. 2016; Menconi et al. 2025). In any case, the strong dependency on light exposure of the *Aft* and *atv* phenotypes highlights the tight relationship existing between anthocyanins and photomorphogenesis in tomato fruits.

Many recent studies have shown the role of light, in particular its intensity, in the biosynthesis of anthocyanins (Ma et al. 2021; Araguirang and Richter 2022; Gonzali et al. 2025). In *Arabidopsis thaliana*, for example, plants grown at higher light intensities accumulate more anthocyanins (Page et al. 2012). The anthocyanin biosynthesis is also regulated by the quality of light: UV and blue light, in particular, have deeper effects compared to other wavelengths, for example red light (Costa Galvão and Fankhauser 2015; Rai et al. 2021). ELONGATED HYPOCOTYL 5 (HY5) and CONSTITUTIVE PHOTOMORPHOGENIC 1 (COP1) represent, respectively, the master positive and negative regulators of photomorphogenesis, acting downstream of the photoreceptors (Jarillo and Cashmore 1998; Teixeira 2020). HY5 is a bZIP TF targeting hundreds of genes through direct binding to their promoters, controlling in this way their expression and thus mediating and transducing the light signals to activate several essential photomorphogenic pathways (Oyama et al. 1997; Gangappa and Botto 2016). HY5 also shows transcriptional and post translational regulations: its transcripts and protein levels directly correlate with light exposure resulting in a regulatory cascade involving light-mediated genes (Ang et al. 1998). Through direct binding to the promoters, HY5 can also control structural and regulatory genes of the flavonoid pathway (Shin et al. 2013; Liu et al. 2018). COP1 is an E3 ubiquitin ligase acting inside a multiprotein complex composed of SUPPRESSOR OF PHYA-105 (SPA), CULLIN 4 (CUL4) and DAMAGE-SPECIFIC DNA BINDING PROTEIN 1 (DDB1) factors (Zhu et al. 2008). The COP1 complex interacts with the CDD complex, containing the proteins DE-ETIOLATED 1 (DET1), DDB1, CONSTITUTIVELY PHOTOMORPHOGENIC 10 (COP10) and CUL4, which acts as an E2 ubiquitin ligase recruiting COP1 targets (Yanagawa et al. 2004). The COP1 multiprotein network represents in plants the main negative regulator of light signalling and photomorphogenesis, targeting, under dark, most of the light-induced factors, including the same HY5 protein, to the 26S proteasome-mediated degradation (Osterlund et al. 1999).

In tomato, *high pigment 1* (*hp1*) and *high pigment 2* (*hp2*) are peculiar mutants known for their phenotypes similar to *COP1* knockout (KO) Arabidopsis plants, except they are not lethal like them (Mustilli et al. 1999; Stacey et al. 2000; Liu et al. 2004; Oravecz et al. 2006). *hp1* and *hp2* mutations were identified in two genes coding for components of the COP1 and CDD complexes, DDB1 and DET1, respectively: the two mutants produce nonfunctional forms of the relative proteins, both resulting in final deactivation of the COP1 network. This leads to stabilization of HY5 and upregulation of light-induced factors and processes, including carotenoid, chlorophyll and anthocyanin synthesis (Liu et al. 2004; Sestari et al. 2014), confirming what already observed in Arabidopsis, where *HY5* overexpression and/or HY5 stabilization both result in higher anthocyanin content under light (Burko et al. 2020).

Although the state of the art pointed out HY5 as master regulator of light-mediated responses and light-triggered anthocyanin synthesis, its silencing, both in *A. thaliana* and in *S. lycopersicum*, surprisingly resulted in plants still able to synthesize anthocyanins under the light stimulus (Holm et al. 2002; Qiu et al. 2019; Han et al. 2020). *Aft/atv/hy5* tomato fruits can in fact produce few anthocyanins in the peel in a light-dependent manner and this phenotype hinted to unknown light-induced and HY5-independent factors, potentially representing the missing part of the light-induced anthocyanin biosynthetic model (Liu et al. 2018; Qiu et al. 2019; Sun et al. 2020).

The quest for the light-induced HY5-independent factors recently came across an important class of TFs, the B-box (BBX) protein family. BBX proteins are a subclass of Zinc-finger TFs characterised by the presence of one or two Zinc-finger B-Box domains, involved in protein–protein interactions (Gangappa and Botto 2014; Yadav et al. 2020). In plants, the B-Box domain is either found alone or together with the “CONSTANS, CO-like, and TOC1” (CCT) domain: BBX proteins are thus classified into five structure groups, according to the number of BBX and CCT domains (Yadukrishnan et al. 2018). It has recently been shown that BBXs can act by the direct or indirect interaction with components of the light signal transduction network, including HY5 and COP1 (Bursch et al. 2020; Luo et al. 2021), thus representing additional regulators of photomorphogenesis, either in a positive or negative way. The most recent models outline the importance of BBX proteins as rate-limiting cofactors of HY5 or as HY5-independent regulators of anthocyanin synthesis (Bursch et al. 2020). They are also negatively regulated by COP1 and are able to induce, directly or indirectly, the synthesis of components of the MBW complex and the expression of some anthocyanin biosynthetic genes (Xu et al. 2016; Bursch et al. 2021; Yang et al. 2022).

In tomato, 29 *BBX* genes were identified (Chu et al. 2016; Lira et al. 2020). Recent studies have shown that the tomato protein encoded by the gene *Solyc01g110180* and named *BBX20*, but corresponding to *BBX25* in Chu et al. (2016) and Lira et al. (2020), can regulate carotenoid and anthocyanin biosynthesis: its overexpression indeed resulted in plants with dark green leaves and fruits (Xiong et al. 2019; Luo et al. 2021). BBX20, which can interact with DET1 and is subjected to COP9-regulated degradation, was shown to regulate the expression of *DFR*, one of the key structural genes involved in anthocyanin biosynthesis, by binding to its promoter (Luo et al. 2021), as well as the expression of the first committed gene in carotenoid biosynthesis (Xiong et al. 2019). More recently, tomato BBX24 was shown to regulate anthocyanin synthesis as well, by interacting either with HY5 or with the components of the MBW complex. Furthermore, *BBX24* silencing in the purple cultivar “Indigo Rose” resulted in retarded anthocyanin coloration of the fruit peel (He et al. 2023). However, due to the intricate patterns shown by the variety of BBX proteins involved in photomorphogenic responses in Arabidopsis plants, it is likely that also in tomato additional BBX factors affect anthocyanin synthesis in tight connection with HY5.

In the present work, two novel tomato BBX proteins were studied in *Aft*/*atv* purple tomato fruits. The two factors resulted to be under light-dependent control and their overexpression led to plants with dark green leaves and intense to deep purple fruits with increased anthocyanin content. The positive effect of these BBXs on the anthocyanin pathway was at different levels: they could activate the expression of *HY5* and stabilize its protein, and they could upregulate anthocyanin structural and regulatory genes, also through physical interaction and additional activation of the MBW complex. In *Aft/atv/hy5* plants, the strong upregulation of one of these *BBX* genes under light well correlated with the residual activation of the anthocyanin pathway in the fruit peel. This work thus highlights the possible role of novel BBXs supporting HY5 in the activation of the anthocyanin synthesis in purple tomato fruits under light.

## Results

### In the *hp2* mutant an increased production of anthocyanins in fruits is accompanied by an alteration of the COP1 network

To study the light regulation of anthocyanin biosynthesis in tomato fruits we used the genotypes *Aft*/*atv* and *Aft*/*atv*/*hp2* (Liu et al. 2004; Sestari et al. 2014). As expected, in *Aft/atv* plants the accumulation of anthocyanins in the fruit peel strongly depended on light intensity, being the content of pigments in the parts of the fruit directly exposed to light higher than that in the shaded parts (Fig. 1A). In the *Aft*/*atv*/*hp2* fruits, on the contrary, anthocyanin synthesis in the epicarp led to a very uniform pigmentation (Fig. 1A). These fruits also displayed a 2- to 4-fold increase in the peel anthocyanin content compared to the *Aft/atv* fruits (Fig. 1A). The expression levels of some biosynthetic and regulatory genes of the anthocyanin pathway (Supplementary Fig. S1) were measured and compared between the lines. The high pigment content of the *Aft/atv/hp2* fruits was due to a strong upregulation of most of the pathway, as shown by the transcription levels of both regulatory and biosynthetic genes. In particular, the genes encoding the three components of the MBW complex activating the anthocyanin biosynthesis, *AN2like,* encoding the R2R3-MYB factor, *AN11*, encoding the WDR protein, and *AN1*, coding for the bHLH factor, were more expressed in *Aft/atv/hp2* than in *Aft/atv* fruits, and the same occurred for the key late biosynthetic structural genes *Dihydroflavonol 4-Reductase (DFR*) and *Anthocyanidin Synthase* (*ANS*) (Fig. 1B).

**Figure 1.**
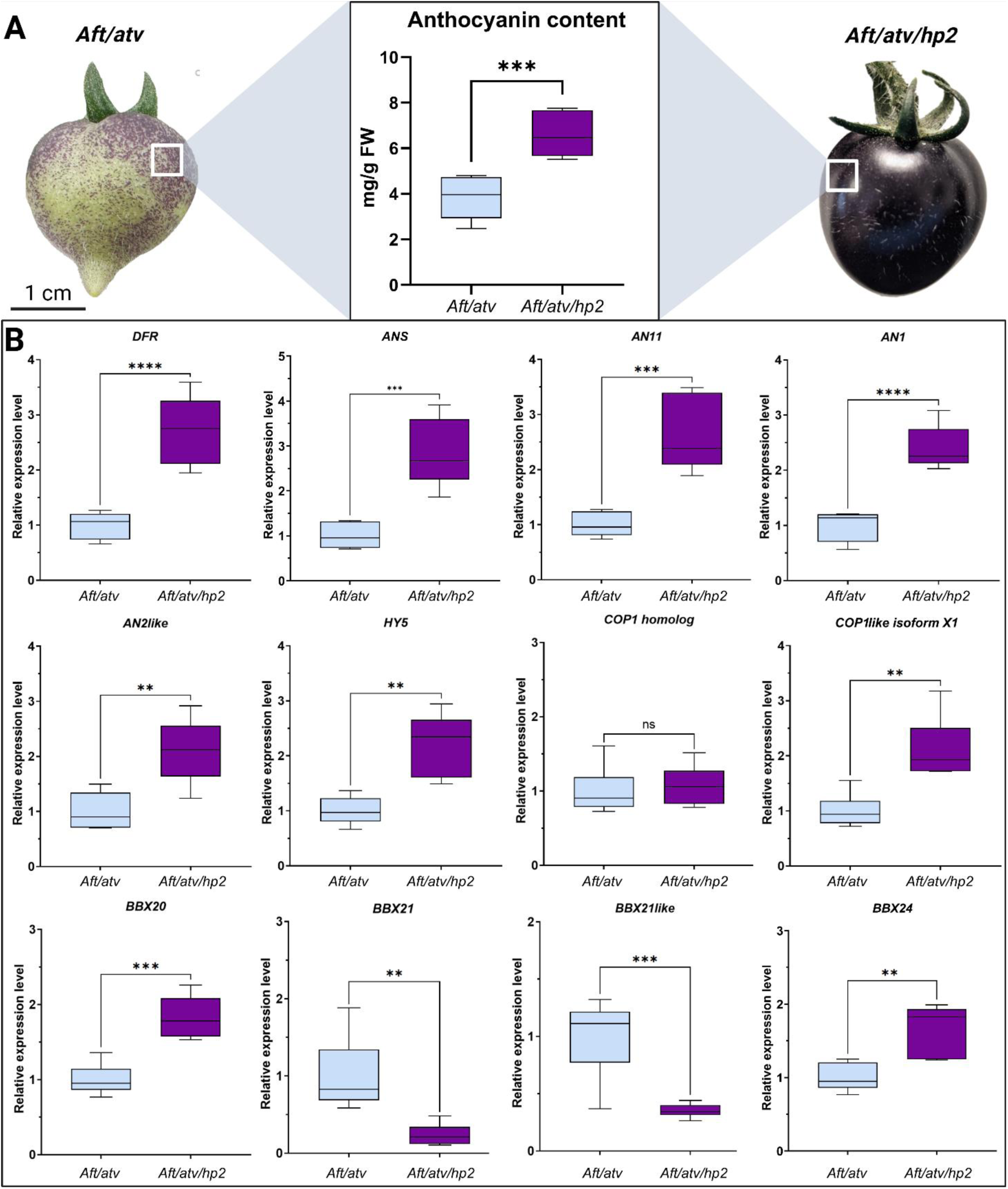
The *hp2* mutation confers upregulation of light-related factors and anthocyanin synthesis. (A) Fruits of *Aft*/*atv* and *Aft*/*atv*/*hp2* lines at the mature green stage and relative anthocyanin quantification in the respective fruit peel. Anthocyanins are expressed as mg petunidin-3-(p-coumaroyl-rutinoside)-5-glucoside g^−1^ fresh weight (FW). Data are means of six biological replicates ± SD. t-test was carried out and **** asterisks indicate significant difference (p≤ 0.0001). (B) Expression analysis of biosynthetic (*DFR*, *ANS*) and regulatory (*AN11*, *AN1*, *AN2like*) genes of anthocyanin synthesis and regulators of photomorphogenesis (*HY5*, *COP1 homolog*, *COP1like isoform X1*, *BBX20*, *BBX21*, *BBX21like*, *BBX24*) performed in the light exposed epicarp of *Aft*/*atv* and *Aft*/*atv*/*hp2* fruits at the mature green stage. Expression levels are shown as relative units, with the value of *Aft*/*atv* set to one. Data are means of six biological replicates ± SD. For each gene expression analysis, t-test was performed between *Aft*/*atv* and *Aft*/*atv*/*hp2* values. Statistical significance is reported in function of the number of asterisks: * p≤0.05, ** p≤0.01, *** p≤0.001, **** p≤0.0001.

The upregulation trend observed in the expression of the anthocyanin genes in the *Aft/atv/hp2* fruits has to be firstly ascribed to the presence of the mutation *hp2* which is known to induce HY5 stabilization (Cañibano et al. 2021; Menconi et al. 2025), with the possible induction of its targets. In confirmation of this, the photomorphogenic genes *HY5* and *COP1like isoform X1*, both HY5 targets (Menconi et al. 2025), resulted more expressed in *Aft/atv/hp2* fruits (Fig. 1B).

The HY5-COP1 network may contribute to the light-dependent regulation of the anthocyanin pathway in a very complex way. We analysed some genes encoding BBX factors, known to often interplay with this network. *BBX20* and *BBX24* are anthocyanin regulators in tomato (Luo et al. 2021; He et al. 2023), and both resulted induced in the presence of the *hp2* mutation (Fig. 1B). Two additional *BBX* genes were then selected, because of the high phylogenetic similarity of their products with the Arabidopsis factor AtBBX21 (Supplementary Fig. S2) which is involved in anthocyanin synthesis in this species (Bursch et al. 2020; Yadav et al. 2020; Yadukrishnan et al. 2018). The two tomato genes were named *BBX21* and *BBX21like* being their proteins both highly similar with AtBBX21 (Supplementary Fig. S2). Interestingly, *BBX21* and *BBX21like* in the *Aft/atv/hp2* line showed a clear reduction in their expression levels compared to the *Aft/atv* one (Fig. 1B), showing an opposite trend of *BBX20* and *BBX24*.

### Structure, cellular localization and activity of the two novel BBX proteins

Tomato BBX21 and BBX21like belong to the structure group IV of the BBX protein family (Lira et al. 2020) and display the typical BOX1 and BOX2 motifs at the N-terminal region, which show high homology with the BOX1 and BOX2 motifs of the Arabidopsis counterpart, while the C-terminal parts, not containing the CCT domain, appear more variable (Fig. 2A).

**Figure 2.**
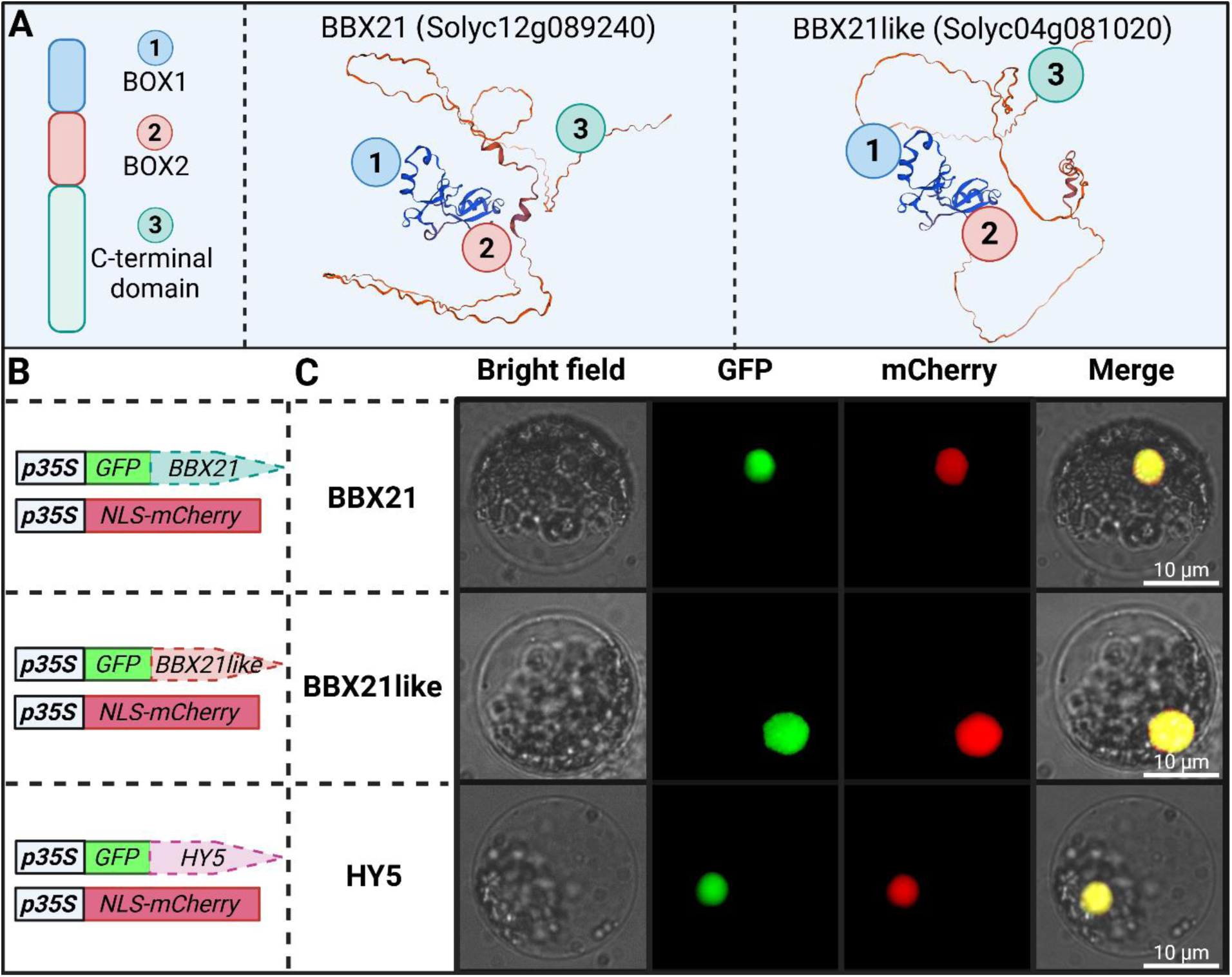
Structural analysis and cellular localization of the BBX factors. (A) Structural modelling of the proteins derived from the genes *BBX21* and *BBX21like* with the typical domains of the BBX factors highlighted. (B) Schematic representation of the genetic constructs used in the localization assays in tomato protoplasts. (C) Cellular localization of the BBX factors and of HY5, after transformation of tomato protoplasts with the corresponding constructs described in panel B. In each transformation, GFP-tagged BBX or HY5 fusion proteins were co-expressed with the Nuclear-Localization Signal (NLS)-mCherry protein. For each localization study, bright field, GFP, mCherry, and merged GFP + mCherry + bright field images of protoplasts are shown.

The cellular localization of the BBX factors was verified in tomato protoplasts, transiently expressing the NLS-mCherry nuclear marker together with the GFP-tagged versions of BBX21 or BBX21like (Fig. 2B). Protoplasts were also transformed to express HY5 (Fig. 2B), which in Arabidopsis interacts with AtBBX21 (Bursch et al. 2020) and is nuclear-localized (Oyama et al. 1997). As expected, also in tomato the two BBX proteins and HY5 localized in the nucleus (Fig. 2C).

To assess their function, BBX21 and BBX21like were evaluated for their ability to regulate the expression of different genes involved in the anthocyanin pathway: *Chalcone Synthase 1* (*CHS1*), *Chalcone Synthase 2* (*CHS2*) and *Flavonoid 3-Hydroxylase* (*F3H*), three EBGs encoding enzymes of the flavonoid pathway; *DFR*, the LBG coding for the enzyme catalysing the first committed step in the anthocyanin synthesis; and the regulatory genes *AN1* and *AN2like* (Supplementary Fig. S1). The sequences of the promoters of all these genes displayed indeed several different BBX and HY5 binding sites (Supplementary Fig. S3). These promoter sequences, driving the firefly *luciferase* (*luc*) reporter, were thus transfected in protoplasts together with effector constructs containing the *BBX21* or *BBX21like* coding sequence (cds) under control of the *35S* promoter (Fig. 3A). The two BBX proteins had an overall positive effect on the activation of all the promoters evaluated (Fig. 3, B-G). The same experiment was repeated with the promoter of *HY5* and also in this case BBX21 and BBX21like activated the reporter gene expression, here up to 10-fold (Fig. 3H).

**Figure 3.**
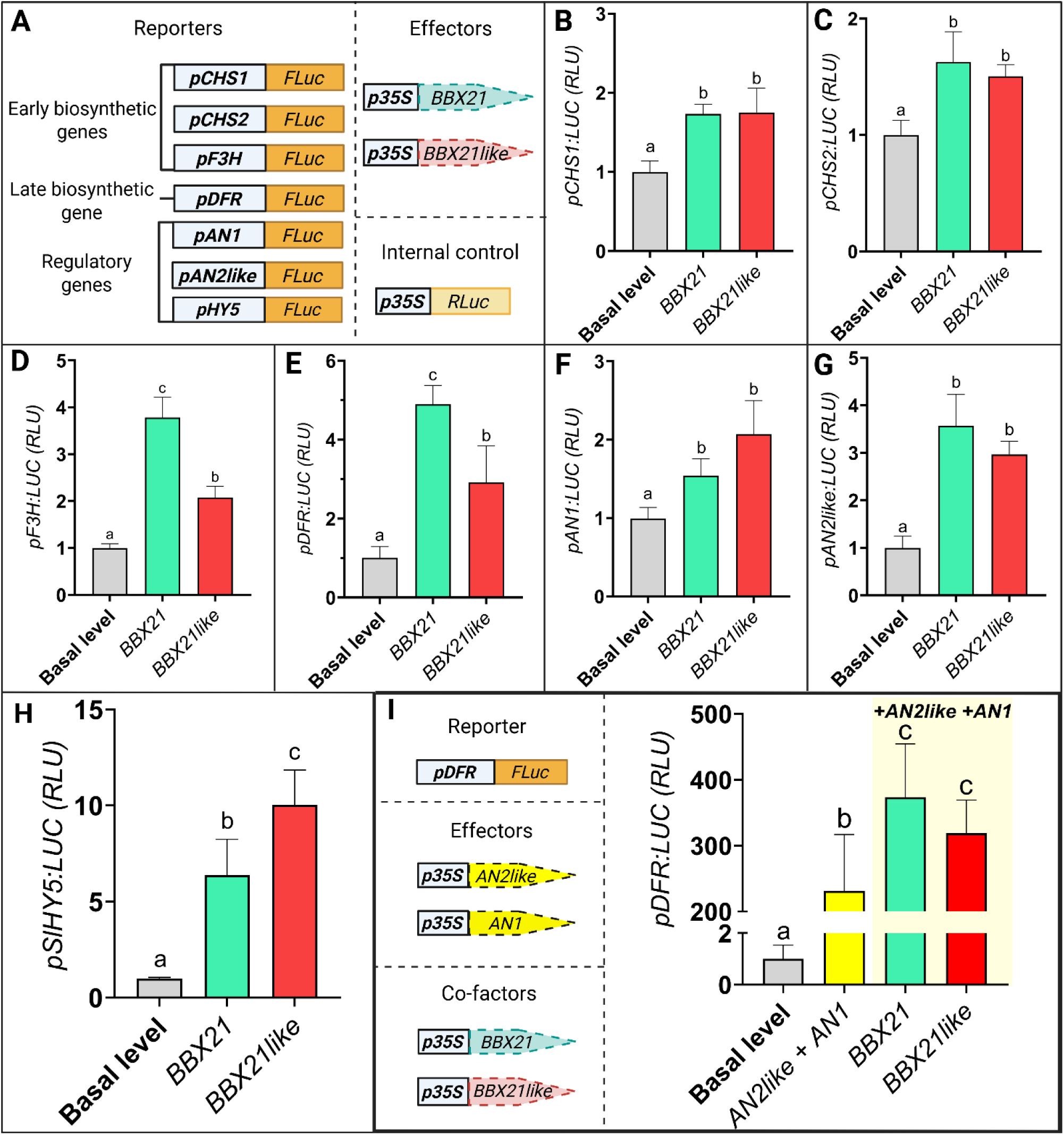
Dual-luciferase assays of the transcriptional activation of anthocyanin genes by BBX21 and BBX21like in tomato protoplasts. (A) Schematic representation of the genetic constructs used in the dual-luciferase assays shown in panels B-H. In the reporter constructs, the promoter of each of the different anthocyanin biosynthetic or regulatory gene drives the firefly *luciferase* (*FLuc*) reporter gene. In the effector constructs, the *p35S* promoter drives the coding sequence (cds) of each of the two BBX encoding genes. The internal control is represented by the *p35S* promoter driving the Renilla *luciferase* (*RLuc*) gene. Transactivation assays of *CHS1* (B), *CHS2* (C), *F3H* (D), *DFR* (E), *AN2like* (F), *AN1* (G) and *HY5* (H) promoters driving *FLuc* gene by BBX21 and BBX21like proteins. Data are expressed as relative luciferase activity (RLU=FLuc/RLuc) with the basal value of the promoters’ luminescence set to 1 and are means of four biological replicates ± SD. In each panel B-H one-way ANOVA with Tukey’s HSD post hoc test was performed. Different letters indicate significant differences at P ≤ 0.05. (I) On the left, schematic representation of the genetic constructs used in the dual-luciferase assay shown in the panel. In the reporter construct, the promoter of the biosynthetic gene *DFR* drives the *FLuc* gene. In the effector constructs the *p35S* promoter drives the expression of the genes encoding the components AN2like and AN1 of the MBW complex; the co-factor constructs are made of the same *p35S* promoter driving each of the two BBX cds. The internal control is represented by the *p35S* promoter driving the *RLuc* gene. On the right, transactivation assay of the promoter of *DFR* driving *FLuc* by AN2like and AN1 co-expressed alone or in the presence of BBX21 or BBX21like. Data are expressed as RLU with the basal value of the *DFR* promoter luminescence set to 1 and are means of four biological replicates ± SD. One-way ANOVA with Tukey’s HSD post hoc test was performed. Different letters indicate significant differences at P ≤ 0.05.

Next, the BBX genes were tested for their ability to affect the activity of the promoter of *DFR*, when activated by the MBW complex. The *DFR* promoter driving the firefly *luc* gene was transfected in protoplasts with *BBX21* or *BBX21like* in the presence of *AN2like* and *AN1* used as effectors (Fig. 3I). AN2like and AN1, expressed together, transactivated the *DFR* promoter with great efficiency, as already known (Colanero et al. 2019). Remarkably, the co-expression of *BBX21* or *BBX21like* resulted in a much greater activation, of at least 30-50% more, of the *DFR* promoter (Fig. 3I).

To confirm the previous results through an independent technique, we performed a Yeast One Hybrid (Y1H) assay, in which the BBX proteins, fused with the *B42* activation domain and expressed under the *GAL1* promoter, were tested for their capacity to bind and activate the expression of promoters of anthocyanin genes and light-regulated genes driving the *GFP* reporter (Fig. 4A; Supplementary Fig. S4). In the Y1H assay BBX21 could bind to the promoters of *CHS1*, *CHS2, F3H, DFR, AN2like, AN1, BBX21,* and *HY5*, with efficiencies ranging from 20- to 400-fold (Fig. 4, B and C). BBX21like interacted with the same promoters as before and also with its own promoter. HY5, putative cofactor of BBX21 in Arabidopsis (Bursch et al. 2020), could also interact with all the promoters analysed, with the exception of *CHS2* (Fig. 4, B and C).

**Figure 4.**
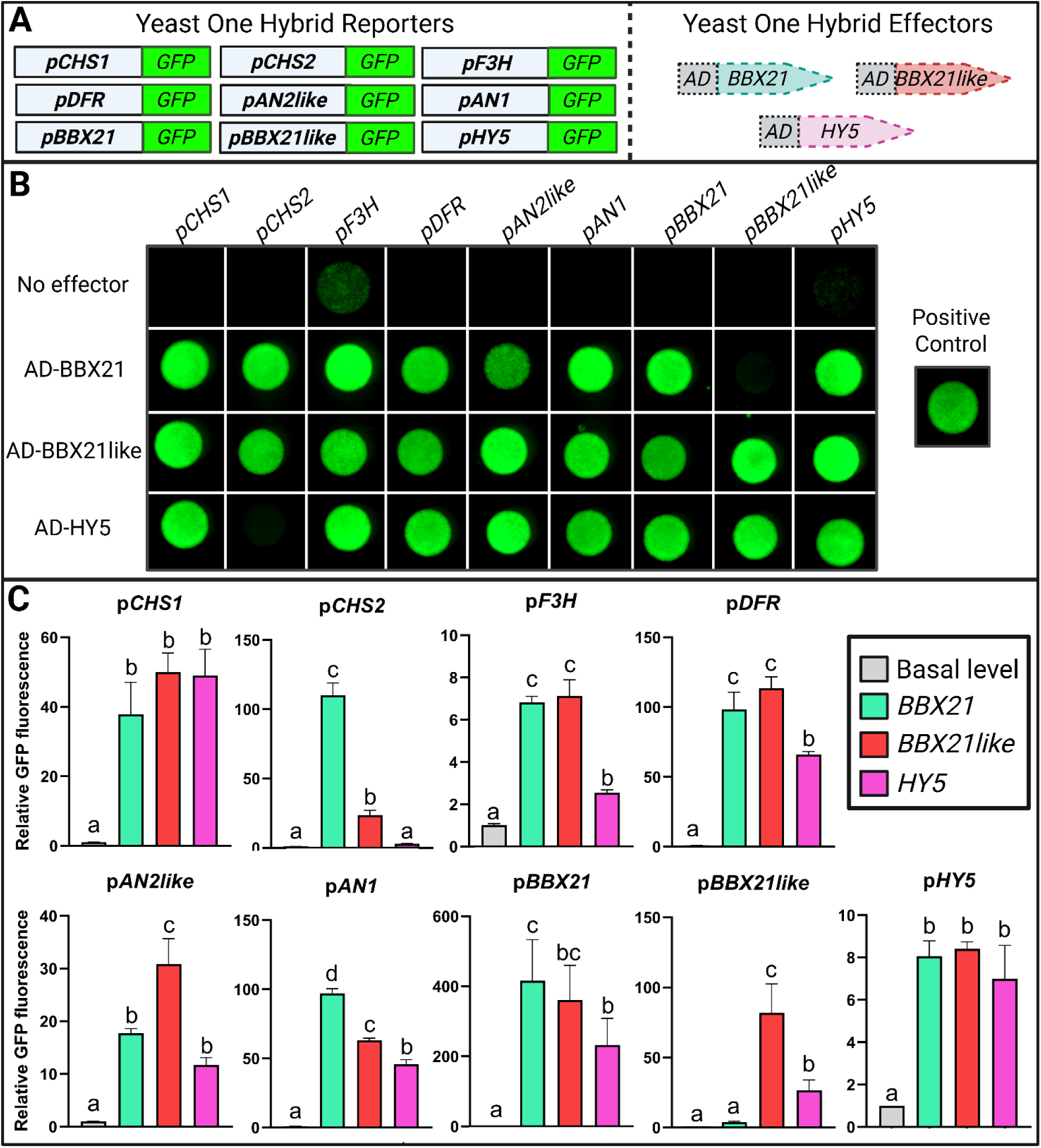
Yeast One Hybrid assay showing the ability of BBX21 and BBX21like to transactivate promoters of anthocyanin and light-related genes. (A) Schematic representation of the genetic constructs used in the Yeast One Hybrid (Y1H) assays. In the reporter constructs the promoter of different anthocyanin (*CHS1*, *CHS2*, *F3H*, *DFR*, *AN2like*, *AN1*) or light signalling (*BBX21*, *BBX21like*, *HY5*) genes drives the *Green Fluorescent Protein* (*GFP*) reporter gene. In the effector constructs the B42 Activation Domain (AD) is fused to the cds of BBX21 or BB21like or of HY5 encoding genes. (B) EGY48 yeast strain transformed with the specified reporter and effector constructs and grown on medium lacking uracile (SD/-U) (No effector) or medium lacking uracile and tryptophan (SD/-U/-T) (bait/pray combinations and positive control). Yeast colonies were screened under UV light. Yeast transformed with only reporter constructs (No effector) were used as negative controls. (C) Quantitative Y1H assays performed on the transformed yeast colonies grown in liquid SD/-U/-T or SD/-U media and quantified for GFP level. Data are expressed as relative GFP levels with the basal value of the reporter set to 1 and are means of three biological replicates ± SD. One-way ANOVA with Tukey’s HSD post hoc test was performed. Different letters indicate significant differences at P ≤ 0.05. Reporter growth controls under white light are shown in Supplementary Fig. S4.

### The stable overexpression of *BBX21* or *BBX21like* increases the anthocyanin content in tomato fruits

Stable transgenic tomato plants overexpressing (OE) *BBX21* or *BBX21like* under the constitutive promoter *35S* were generated in the *Aft*/*atv* background. Three independent transgenic lines were produced for each *BBX* gene. The *BBX21-* and *BBX21like-*OE plants resulted in thick dark green leaves (Supplementary Fig. S5) and short stems (Supplementary Fig. S6) and produced intense to deep purple fruits (Fig. 5, A-B). The anthocyanin quantification performed in the fruit peels confirmed an increase in the anthocyanin content compared to the *Aft*/*atv* background, ranging from 2- to 4-fold (Fig. 5C). Noteworthy, whereas the upregulation of *BBX21* had no effect on the *BBX21like* transcript level, the *BBX21like-*OE lines showed also upregulation of *BBX21* (Fig. 5D). The lines with the highest levels of transgene expression and anthocyanins in fruits (Fig. 5), corresponding to Line A for *BBX21* and Line D for *BBX21like,* were selected for more detailed characterizations.

**Figure 5.**
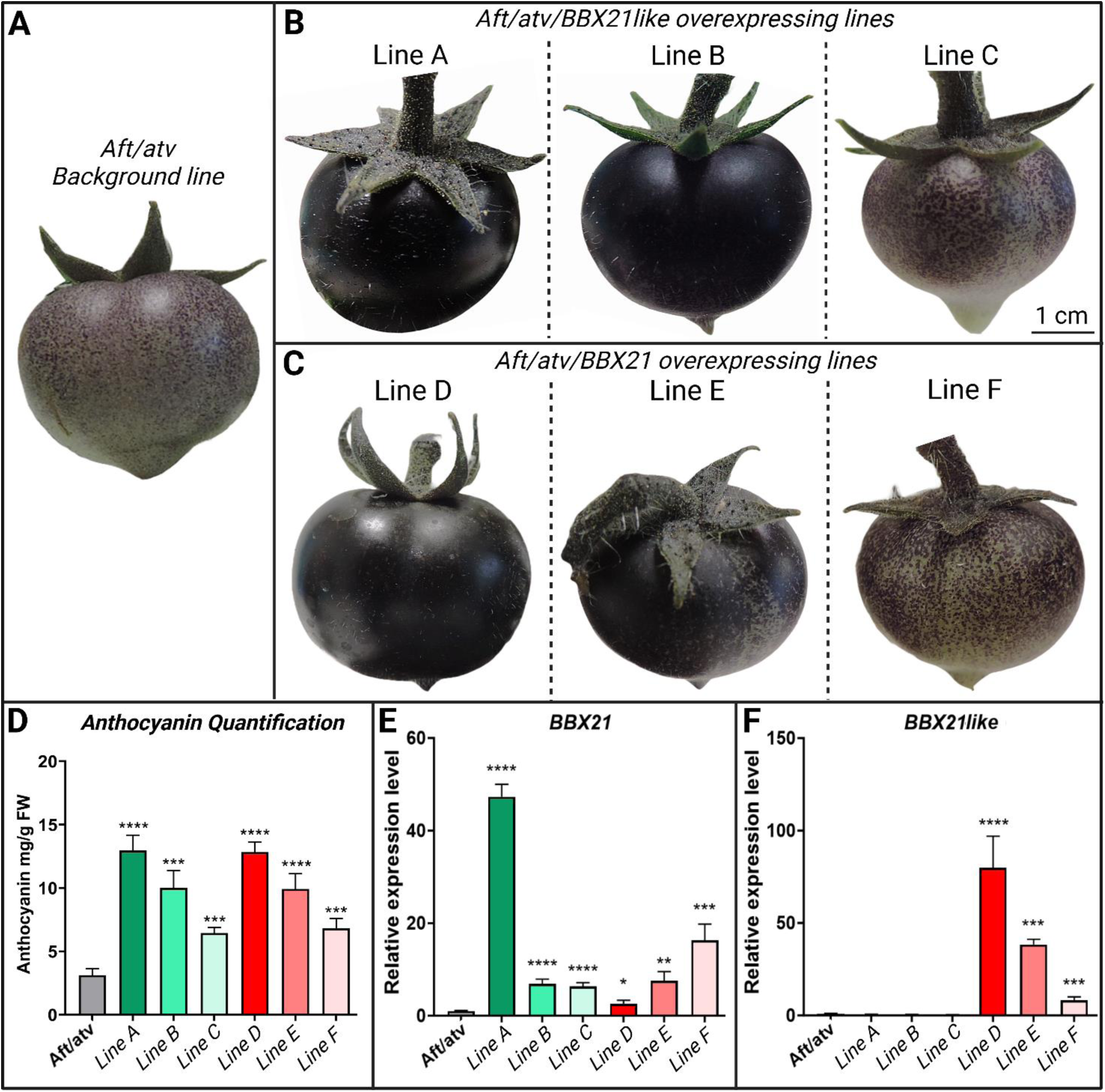
Overexpression of BBX factors in the *Aft*/*atv* MicroTom background. (A-B) Phenotypes of the fruits obtained from three independent lines overexpressing (OE) *BBX21* (Line A, Line B, Line C) or *BBX21like* (Line D, Line E, Line F) coding sequences compared to their *Aft/atv* background. (C) Anthocyanin quantification in the fruit epicarp at the mature green stage of the OE lines. Anthocyanins are expressed as mg petunidin-3-(p-coumaroyl-rutinoside)-5-glucoside g^−1^ fresh weight (FW). Data are means of 10 biological replicates ± SD. Quantitative analysis of the transcript levels of *BBX21* (D) and *BBX21like* (E) in peel from mature green fruits of the OE lines in comparison with the *Aft/atv* background. Expression levels are shown as relative units, with the value of *Aft*/*atv* set to one. Data are means of 10 biological replicates ± SD for each genotype. In panels C-E, t-test was performed comparing each line with the *Aft*/*atv* background. Statistical significance is reported in function of the number of asterisks: * p≤0.05, ** p≤0.01, *** p≤0.001, **** p≤0.0001.

A transcriptomic analysis was carried out in the fruit peel of the transgenic lines. Both *BBX21*- and *BBX21like*-OE lines displayed a clear upregulation of biosynthetic genes involved in the anthocyanin pathway (Fig. 6, A-C and G; Supplementary Fig. S7). *CHS1* resulted the most upregulated *EBG*, reaching an induction level ranging between 5- to 10-fold compared to control. *CHIlike, F3H, DFR* and *ANS* showed increased expression levels as well, ranging in these cases between 2- to 4-fold. *PAL5* and *CHS2* resulted induced only in some lines (Supplementary Fig. S8). The transcription levels of the regulatory genes encoding early activators of the biosynthetic pathway, such as the R2R3-MYB factor MYB12, or components of the MBW complex inducing the late steps of the anthocyanin synthesis, such as AN2like, AN1 and AN11, varied in different ways: *MYB12* resulted upregulated in only two lines and *AN2like* was almost not affected by the overexpression of any of the two BBXs whereas most transgenic lines showed a clear upregulation of *AN1* and *AN11* (Fig. 6, D-G; Supplementary Fig. S8).

**Figure 6.**
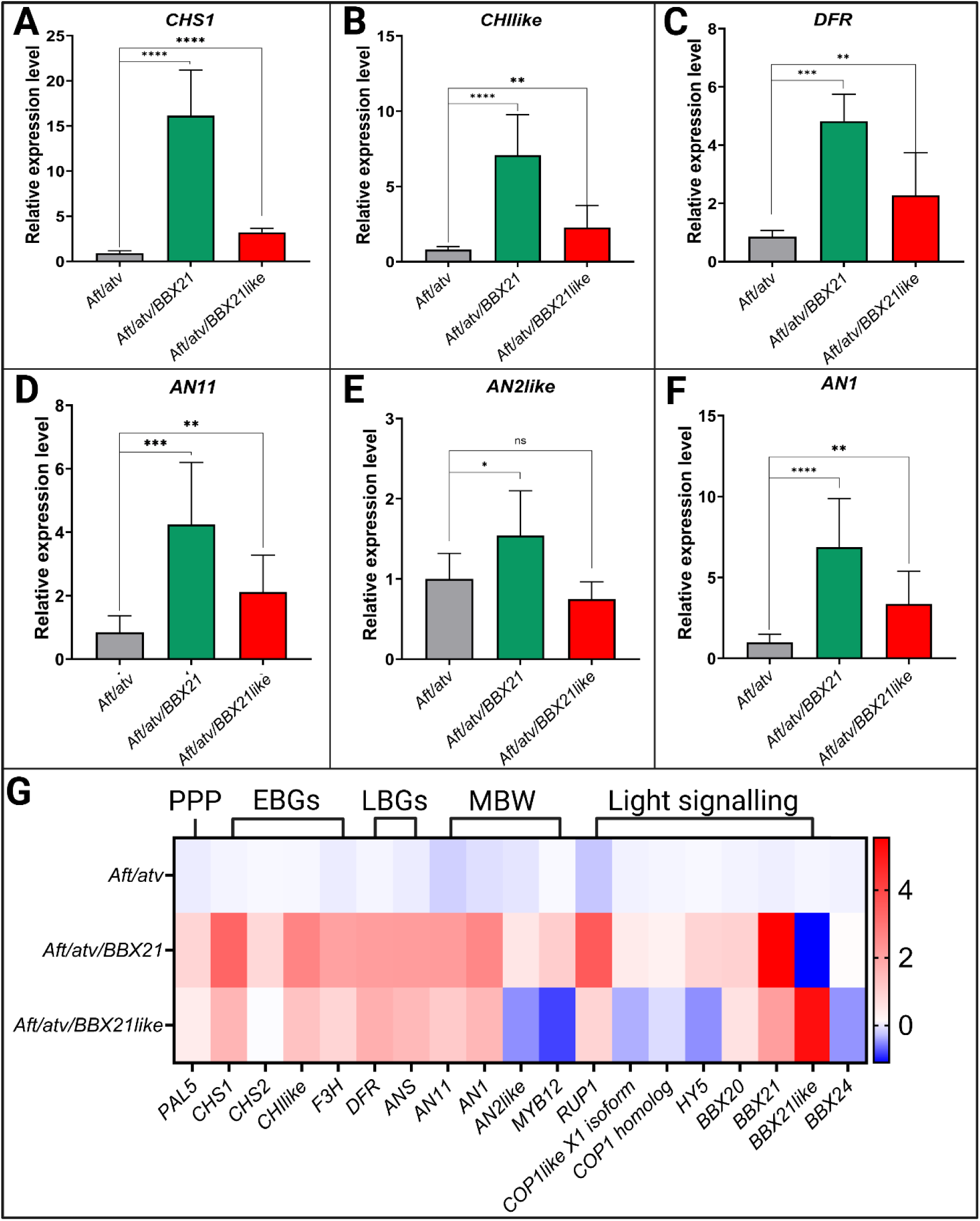
The overexpression of the BBX proteins affects anthocyanin and light-related genes. Quantitative analysis of the transcript levels of some biosynthetic [*CHS1* (A), *CHIlike* (B), *DFR* (C)] and regulatory [*AN2like* (D), *AN1* (E), *AN11* (F)] genes of the anthocyanin pathway in peel from mature green fruits of *BBX21*- and *BBX21like*-overexpressing (OE) lines and of their *Aft*/*atv* background. (G) Heatmap showing the entire set of gene expression data measured in fruits, as above described, containing biosynthetic and regulatory genes involved in anthocyanin synthesis and other photomorphogenic processes. Genes are grouped based on their functions: phenylpropanoid pathway (PPP); anthocyanin early biosynthetic genes (EBGs); anthocyanin late biosynthetic genes (LBGs); MYB, bHLH and WDR components of the MBW complex (MBW) encoding genes and light signalling genes. In A-F, expression levels are shown as relative units, with the value of *Aft*/*atv* set to one. Data are means of 10 biological replicates ± SD for each genotype. t-test was performed comparing each line with background. Statistical significance is reported in function of the number of asterisks: * p≤0.05, ** p≤0.01, *** p≤0.001, **** p≤0.0001. In G, relative expression units for each gene were transformed by Log2(x). Expression levels are represented as a heatmap by the colour and intensity of the boxes according to the colour scale. The corresponding bar plots with statistical analyses for each gene are shown in Supplementary Fig. S7 and Fig. S8.

Genes coding for some components of the light signalling network were analysed (Fig. 6G; Supplementary Fig. S8): compared to control, *COP1 homolog* resulted not affected in the transgenics, whereas *COP1like isoform X1* appeared modulated in a positive way only in the *BBX21-*OE Line A, which contemporarily showed also a 2-fold upregulation of *HY5* (Supplementary Fig. S8). The same line, that was the one producing more anthocyanins, showed again the highest upregulation of almost all the genes analysed. Interestingly, the gene *RUP1* resulted strongly upregulated in most *BBX*-overexpressing fruits. *BBX20* was slightly upregulated in most of the lines and *BBX24* was downregulated in *BBX21like* transgenic lines (Fig. 6G; Supplementary Fig. S8).

### BBX21 and BBX21like physically interact with components of the light signalling machinery and of the anthocyanin synthesis and stabilize HY5

BBX factors were previously shown to participate in protein interactions with other components of the light signalling machinery. A Yeast Two Hybrid (Y2H) assay was thus performed to test putative interactions of BBX21 and BBX21like with anthocyanin regulators (AN2like and AN1) or light signalling-related factors (HY5, DET1, COP1like isoform X1, COP1 homolog) and among them (Fig. 7; Supplementary Fig. S9). BBX21 and BBX21like were able to physically interact with all the tested factors, themselves included, as in the Y2H assay the relative bait/prey combinations allowed transformed yeast cells to grow on selective media and to perform β-galactosidase activity (Fig. 7A).

**Figure 7.**
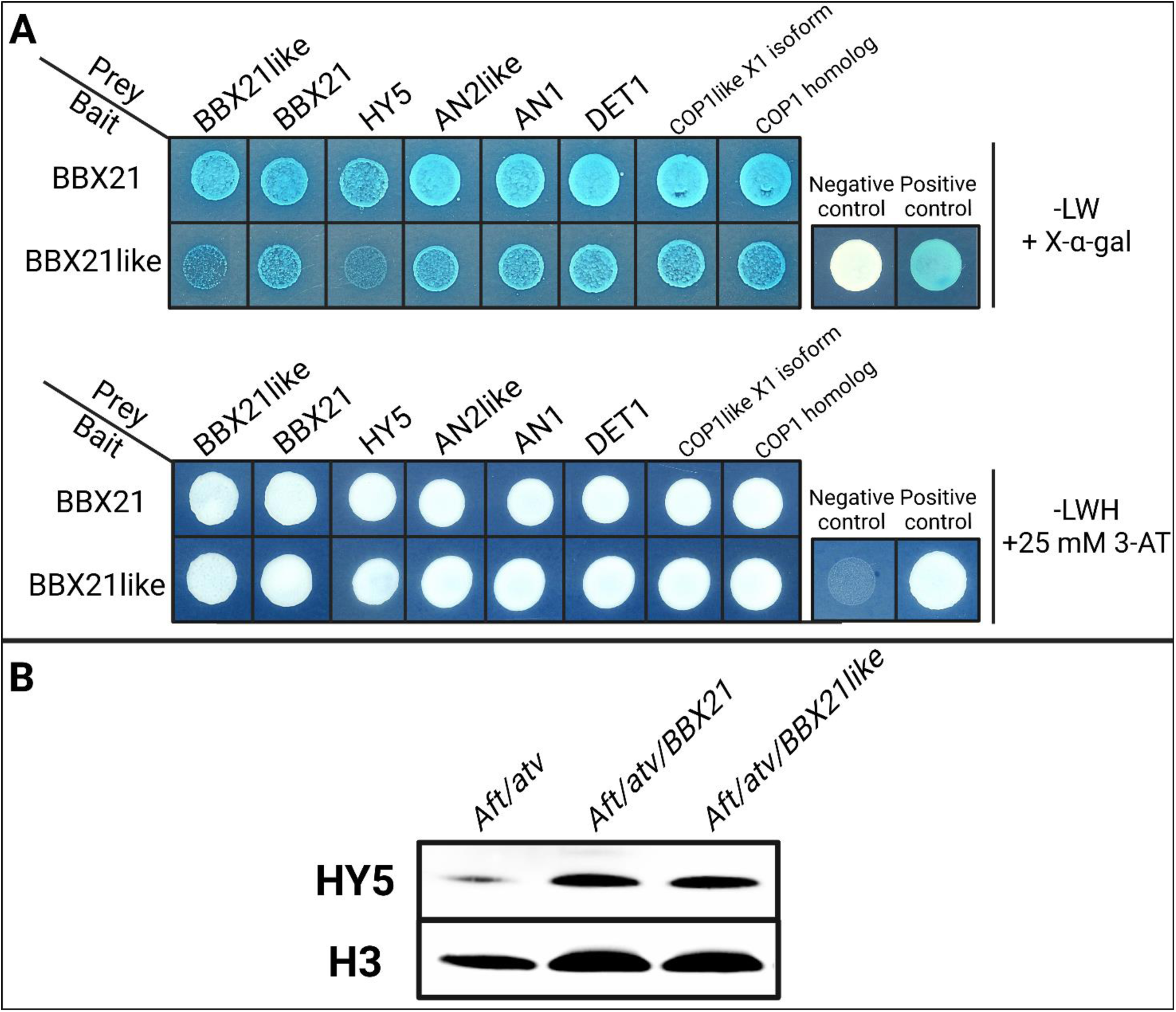
Interaction of the BBX factors with proteins involved in anthocyanin synthesis or light signalling and effects of *BBX* overexpression on HY5 levels. (A) Results of the Yeast Two Hybrid (Y2H) assay: MAV203 yeast strain transformed with the reported bait and prey constructs and grown on medium lacking leucin (L) and tryptophan (W) supplemented with X-α-Gal (-LW + X-α-Gal), and on medium lacking L, W, histidine (H) and supplemented with 25 mM of 3-AT (-LWH + 25 mM 3-AT). The negative controls for the Y2H assay are shown in Supplementary Fig. S9. (B) Representative Western-blot carried out with nuclear proteins isolated from peel of *Aft*/*atv*, *Aft*/*atv*/*BBX21-overexpressing (OE)* and *Aft*/*atv*/*BBX21like-OE* fruits probed with specific antibodies for HY5 and histone H3. The uncropped unedited membranes are shown in Supplementary Fig. S10.

Next, we quantified the protein levels of HY5, known to be stabilized by these classes of BBX factors in Arabidopsis (Podolec et al. 2022). Western blot analysis was carried out on nuclear fractions extracted by either fruit peel or leaf of *Aft/atv* and *BBX*-OE plants. In fruits, overexpression of *BBX21* or *BBX21like* resulted in higher HY5 nuclear levels when compared to the *Aft/atv* background (Fig. 7B; Supplementary Fig. S10). In leaves, the increase in the HY5 protein levels was visibly lower than in fruits (Supplementary Fig. S11).

### Knockout plants for *HY5* produce less anthocyanins in fruits and only in later ripening stages

Stable KO plants for *HY5* were previously produced through CRISPR/Cas9 technology in the *Aft*/*atv* background (Menconi et al. 2025). *Aft/atv/hy5* plants displayed elongated stems (Supplementary Fig. S12) and produced fruits with drastically lower amounts of anthocyanins in the fruit epicarp compared to control (Fig. 8, A-D). Moreover, *Aft/atv/hy5* fruits developed slower than control fruits and at the mature green (MG) stage it was possible to distinguish two different phases of pigmentation. In the *Aft/atv* line anthocyanins started to be synthesized at the onset of the MG stage (early MG) and then continued to accumulate till the late MG stage (late MG) (Fig. 8, A and B). In the *Aft/atv/hy5* line, anthocyanin synthesis was delayed, with fruits showing complete absence of pigments at early MG, being completely pale green, whereas at late MG they showed a moderate anthocyanin accumulation in the light-exposed parts of the epicarp (Fig. 8, C and D). However, the anthocyanin content in *Aft/atv/hy5* late MG fruits was much lower than that in their *Aft/atv* counterparts, and it was even lower than the one measured in *Aft/atv* early MG fruits (Fig. 8E).

**Figure 8.**
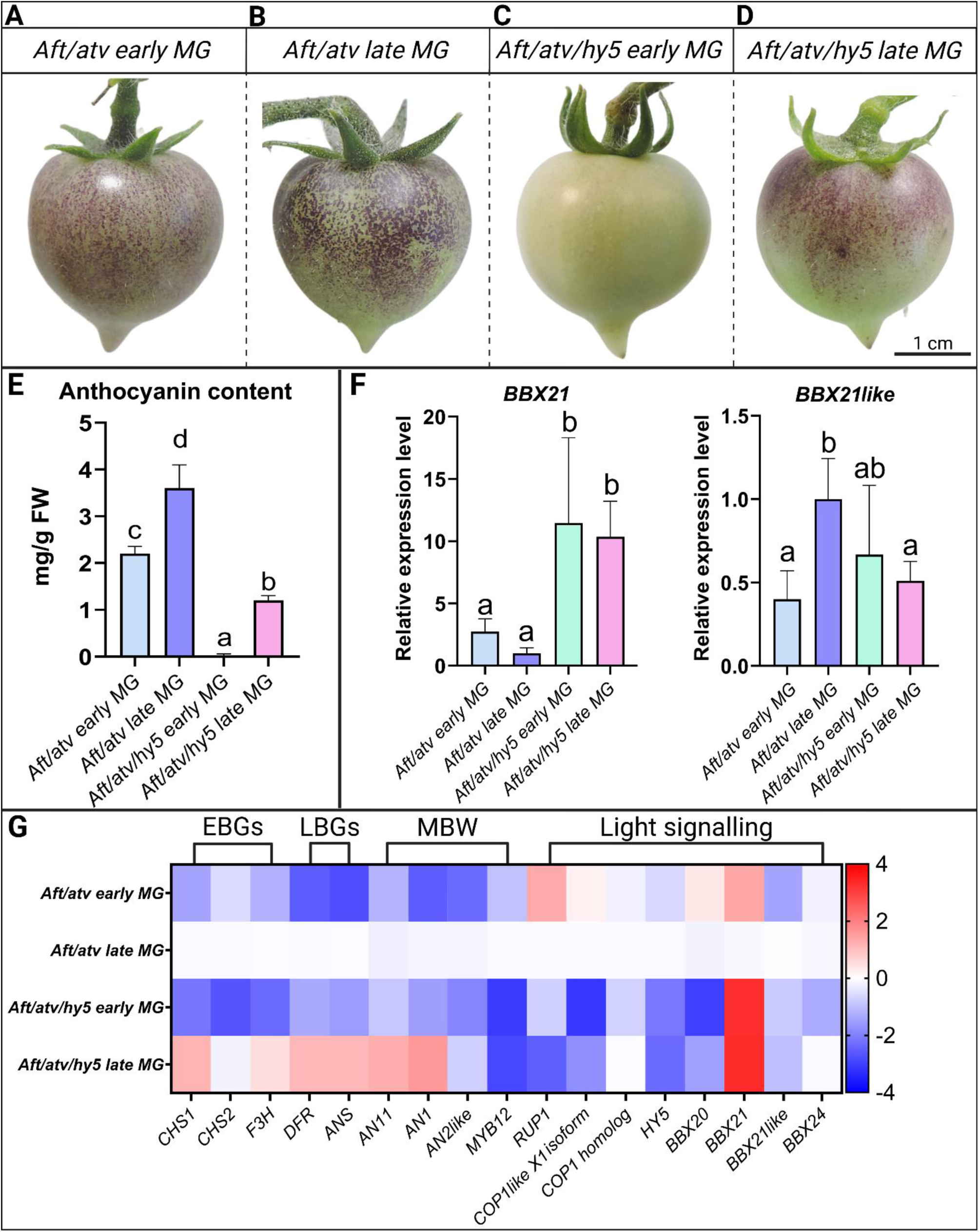
Loss of HY5 leads to strong repression of anthocyanin synthesis and deregulation of *BBX21* expression. (A-D) Phenotypes of the fruits obtained from the *Aft*/*atv* and *Aft*/*atv*/*hy5* lines at early and late mature green (MG) stages. (E) Anthocyanin quantification in the fruit epicarp of the *Aft*/*atv* and *Aft*/*atv*/*hy5* lines at early and late MG. Anthocyanins are expressed as mg petunidin-3-(p-coumaroyl-rutinoside)-5-glucoside g^−1^ fresh weight (FW). Data are means of 6 biological replicates ± SD. (F) Quantitative analysis of the transcript levels of *BBX21* and *BBX21like* in fruit peel at early and late MG of the *Aft*/*atv*/*hy5* line compared with the *Aft*/*atv* background. Expression levels are shown as relative units, with the value of *Aft*/*atv* at the late MG stage set to one. Data are means of 6 biological replicates ± SD for each genotype. In panels E-F one-way ANOVA with Tukey’s HSD post hoc test was performed. Different letters indicate significant differences at P ≤ 0.05. (G) Heatmap showing the entire set of gene expression data measured in peel from early and late MG fruits of *Aft*/*atv*/*hy5* compared with *Aft*/*atv* background. Genes are grouped based on their functions: phenylpropanoid pathway (PPP); anthocyanin early biosynthetic genes (EBGs); anthocyanin late biosynthetic genes (LBGs); MYB, bHLH and WDR components of the MBW complex (MBW) encoding genes and light signalling genes. In G, relative expression units for each gene were transformed by Log2(x). Expression levels are represented as a heatmap by the colour and intensity of the boxes according to the colour scale. The corresponding bar plots with statistical analyses for each gene are shown in Supplementary Fig. S13.

A transcriptomic analysis was carried out in the fruit peel of the two different lines (Fig. 8, F and G; Supplementary Fig. S13). The early MG stages of either *Aft/atv* or *Aft/atv/hy5* fruits were characterized by lower expressions of early (*CHS1, CHS2, F3H*) and late (*DFR, ANS*) biosynthetic genes as well as of regulatory genes (*AN11, AN1, AN2like*) compared with the respective late MG stages, well correlating with the anthocyanin contents of the respective fruits. Interestingly, in *Aft/atv/hy5* fruits *AN2like*, and even more *MYB12*, remained lowly expressed at late MG.

Genes coding for some components of the light signalling network were analysed as well (Fig. 8G; Supplementary Fig. S13). *HY5* resulted downregulated in the *Aft/atv/hy5* fruits, at both early and late MG stages, and this trend was also shown by *RUP1*, *COP1like X1 isoform* and *BBX20*. On the contrary, *COP1 homolog* and *BBX24* were quite similar in their expression levels between the two genotypes.

Genes coding for the two BBX factors under study were finally analysed (Fig. 8, F and G). *BBX21* was strongly upregulated, up to 10-fold, in the *Aft/atv/hy5* fruits at both MG stages, while *BBX21like* resulted less expressed compared to control fruits at late MG.

## Discussion

In the last years and in different plant species BBX factors have been shown to participate in light signal transduction and anthocyanin synthesis through the HY5-COP1-mediated switch (Xu et al. 2016; Xiong et al. 2019; Cao et al. 2023). In Arabidopsis, the HY5 light signalling pathway resulted tightly interconnected with different BBX proteins, which in part act as cofactors, either positively or negatively modulating HY5 activity, in part control *HY5* at transcriptional level and in part are in turn transcriptionally regulated by HY5 which can bind to their promoters (Cao et al. 2023). In tomato, BBX20 and BBX24 have been recently studied, and both resulted able to positively affect anthocyanin synthesis (Xiong et al. 2019; Luo et al. 2021; He et al. 2023), but the BBX family includes other factors which may be involved in the process. For this reason, the candidate genes *BBX21* and *BBX21like* were selected based on their similarities with the *A. thaliana* BBX21 factor, which is clearly involved in photomorphogenesis and anthocyanin regulation (Xu et al. 2016; Bursch et al. 2020; Cao et al. 2023).

Tomato BBX21 and BBX21like show similar structures and are both closely related to AtBBX21: they could have been originated from a gene duplication event, and the high sequence similarity suggests potential redundancy. Furthermore, BBX21 and BBX21like are also very similar to tomato BBX20, which was recently shown to positively regulate carotenoids and anthocyanins and to be target of the COP9 signalosome (Xiong et al. 2019; Luo et al. 2021). Interestingly, while *BBX20* and *BBX24* resulted both upregulated in the anthocyanin-enriched *Aft/atv/hp2* fruits compared to the *Aft/atv* ones suggesting the existence of a HY5-mediated transcriptional induction, *BBX21* and *BBX21like* resulted both downregulated: thus, their transcription appeared to be negatively affected as consequence of COP1 deactivation and/or HY5 stabilization.

Functional *in vitro* studies indicated that the two novel BBX proteins are nuclear localized, similarly to HY5, and activate the promoters of anthocyanin biosynthetic and regulatory MBW genes, with BBX21 being the most consistent in the effects. On the other hand, in the same assays HY5 resulted able to bind promoters of either anthocyanin biosynthetic or MBW genes, in particular *AN2like*(*R2R3-MYB*) and *AN1* (*bHLH*), in line with studies performed in other species (Stracke et al. 2010; Shin et al. 2013). The mild activations of the anthocyanin gene promoters shown by the BBXs in the protoplast assays, included in a 2- to 4-fold range, are similar to results obtained with other BBXs (Xu et al. 2016; Xiong et al. 2019; Luo et al. 2021), but might imply the necessity of a cofactor, for example the same HY5 (Bursch et al. 2020), to fully perform their activity. Interestingly, BBX21 and BBX21like activated *in vitro* also the promoter of HY5, suggesting that they might reinforce *in planta* the light response mediated by HY5. Moreover, each of the two BBX factors was able to bind to its own promoter and to the promoter of the other *BBX* gene, and the same HY5 could bind to the promoters of *BBX21* and *BBX21like,* indicating the existence of complex regulatory control loops.

*In vitro* transactivation assays and protein-protein interaction studies highlighted the existence of a physical interaction of BBX21 and BBX21like with AN2like and AN1 gene products and a further possible cooperation in the transcriptional induction of the anthocyanin LBGs with the MBW complex containing the same R2R3-MYB and bHLH factors. In this way, higher transcriptional levels of *AN2like* and *AN1*, induced by HY5 and reinforced by BBX21 and BBX21like activities, would first lead to higher MBW complex levels and then to the hypothetical formation of quaternary complexes made of MBW components and BBX factors (MBW-B), potentially showing higher transcriptional activity or increased stability.

Overexpression of *BBX21* and *BBX21like* in *Aft/atv* plants produced deep purple fruits, accumulating anthocyanins independently of direct light exposure, and thick and dark green leaves, as well as shorter stems, phenotypes very similar to those of the *Aft/atv/hp2* plants. This confirmed that BBX21 and BBX21like may act as positive regulators of anthocyanin synthesis and other photomorphogenic responses *in vivo*, with BBX21 showing higher efficiency in activating anthocyanin synthesis than BBX21like.

The transcript analysis carried out in the fruits of the *BBX*-OE plants confirmed that, even if with different effectiveness, the overexpression of each BBX in the *Aft/atv* background led to increased induction of both anthocyanin EBGs and LBGs as well as of some regulatory genes. This would suggest similar functions of the two BBXs as transcriptional activators or, in alternative, since overexpression of *BBX21like* led to upregulation of *BBX21*, a possible convergence among the downstream targets on BBX21 activity, which indeed showed the highest efficiency in activation of the pathway.

Among the regulatory genes, the strongest impulse exerted by BBX21 was on the transcripts of *AN1* and *AN11*, whereas *MYB12*, activating the EBGs (Stracke et al. 2010), appeared only moderately induced, and *AN2like* very slightly affected. The fact that BBX21 and BBX21like appeared to interact stronger with *AN2like* promoter when overexpressed *in vitro* than *in vivo* might suggest that this *R2R3-MYB* gene does not constitute a primary target of their regulation rather representing a preferred target of HY5 under light, as clearly confirmed by the increased transcription of *AN2like* in the presence of the *hp2* mutation. It is likely that the levels of *AN2like* induced by light through HY5 were sufficient to activate at a basal level the anthocyanin pathway, as shown by the moderate pigmentation shown by the *Aft*/*atv* fruits. The overexpression of the BBX factors, leading to upregulation of *AN1* and *AN11,* would increase the amounts of the other two, possibly limiting, components of the MBW complex, with the consequent strengthened transcription of the LBGs observed in the *BBX*-OE lines. The autoactivation of *AN1* exerted by the same MBW complex in a feedforward mechanism (Sun et al. 2020) would further contribute to increase the transcription of the downstream biosynthetic and regulatory genes. All these mechanisms would reinforce the basal activation of the anthocyanin pathway of the *Aft*/*atv* fruits, increasing the anthocyanin pigmentation as seen in the fruits of the *BBX*-OE plants.

*In vivo* the overexpression of the two BBX factors did not influence in a particular way the other light signalling genes, suggesting that their function in the signalling cascade was not as central as the role of HY5. An exception was represented by *RUP1* which codes for a WD40 protein regulating the UV-B response. AtRUP1 is induced by UV light and is positively regulated by HY5; its role is to trigger the dimerization of the UV receptor UVR8, thus preventing the UVR8-induced COP1 repression and the consequent HY5 stabilization: it acts in this way as a negative photomorphogenic regulator (Zhang et al. 2021). The activation of *RUP1* exerted by the two BBX proteins suggests a possible role of these factors in the regulation of the UV-B mediated photomorphogenic response in tomato, similarly to the one carried out by AtBBX21 in Arabidopsis (Podolec et al. 2022).

However, in the complex of the photomorphogenic responses, the physiological role in tomato plants of the BBX factors under study might be more profound than that of single and simple transcriptional activators: they indeed demonstrated to be able to perform homo- or heterodimerizations, to form complexes with the MBW members, as above discussed, and also interact with HY5, DET1 and the two COP1 proteins, these last involved in HY5 destabilization in the dark (Menconi et al. 2025). Moreover, the overexpression of BBX21 and BBX21like increased the protein levels of HY5 in tomato fruits, similarly to what seen in Arabidopsis, where AtBBX21 can post-translationally stabilize HY5 (Podolec et al. 2022). Putting together all these results, one might hypothesize that BBX21 and BBX21like act as shields to HY5 degradation. Being the BBXs preferential targets for COP1 activity, this would decrease the levels of BBX factors in the cell but might confer a cellular status more prone to respond to light stimuli, given the higher levels of HY5.

In Arabidopsis the stabilized HY5 triggers a negative feedback loop, in turn repressing the expression of *BBX21* and *BBX22*, fine-tuning in this way the photomorphogenic responses (Xu et al. 2016). This mechanism appears to be conserved in tomato, given that *BBX21* and *BBX21like* were downregulated in the *Aft/atv/hp2* fruits, and this downregulation might be consequence of the upregulation of HY5 due to the *hp2* mutation.

Stable knockout of *HY5*, with fruits only showing a mild anthocyanin pigmentation, confirmed that HY5 protein is required for a strong anthocyanin accumulation, but also that other TFs can activate the pathway in its absence. While *BBX21like* resulted not much affected by the loss of HY5*, BBX21* was strongly upregulated in *Aft/atv*/*hy5* fruits during the MG stage, and this might be at the base of the residual anthocyanin response in the *hy5* mutant. In *Aft/atv* fruits BBX21 might contribute to activate *HY5* and to stabilize its protein which, at a certain threshold, triggers the repression of the same *BBX21* expression to fine-tune the level of anthocyanins, in a negative feedback loop. The loss of *HY5* might disrupt this mechanism: in *Aft/atv/hy5* fruits *BBX21*, subtracted from the negative feedback exerted by HY5, might be expressed under light at a level sufficient to start an autoactivation process, progressively reinforcing its own transcription and inducing, even if slowly, anthocyanin biosynthesis. Anyway, the anthocyanin synthesis fails to reach the levels observed in *Aft/atv* fruits, probably because HY5 is necessary to fully activate the pathway: *MYB12* and *AN2like*, for example, were consistently less expressed in *Aft/atv*/*hy5* MG fruits than in *Aft/atv* and their low transcriptions might explain the weak activation of EBGs and LBGs, respectively.

## Conclusion

In the present research, we show that tomato BBX21 and BBX21like are light-dependent coregulators of anthocyanin synthesis in purple tomato fruits and that their direct targets can be either the biosynthetic genes or the *WDR* and *bHLH* genes coding for the two corresponding components of the MBW complex. The *R2R3-MYB* gene, encoding the third component of the complex, that has always thought to be the most tuneable, appeared on the contrary to be largely under HY5 control.

BBX21 and BBX21like resulted similar in some activities and thus partially redundant, possibly also for a limited convergence of common secondary targets. Furthermore, they could not only trigger anthocyanin synthesis directly binding to gene promoters but also interacting with the MBW complex further increasing its transcriptional activity. Finally, BBX21 appeared to play a role in shaping anthocyanin synthesis in the absence of *HY5,* partially compensating its loss.

The relative upstream position of the BBX factors towards the anthocyanin pathway, tightly related to light signalling and photomorphogenesis, results in a very complex network, regulating several different genes and proteins through protein and genetic interactions. By putting together literature and results from the present study, the key findings in the light-activation of anthocyanin synthesis in purple tomato fruits can be summarized as in the schemes of Fig. 9 and Fig. 10. HY5 is a major factor positively regulating anthocyanin synthesis under light but also controlling several other proteins, including BBX and MYB factors, with COP1 acting as a major negative regulator. COP1 deactivation due to the *hp2* mutation results in downregulation of *BBX21* and *BBX21like* expressions, probably due to the upregulation of HY5 which represses their transcription. BBX21 and BBX21like act as positive transcriptional and post-translational regulators of HY5 but may also be under its negative transcriptional control, at least when HY5 levels exceed a certain threshold, such as in presence of the *hp2* mutation which subtracts HY5 protein from COP1-mediated degradation. In this sense, the two BBXs would not act as HY5-independent factors but rather as HY5-coregulators, fine-tuning HY5 functions and preventing excessive anthocyanin synthesis, once activated. Only in case of HY5 absence, as in the *Aft/atv/hy5* line, BBX21 would vicariate it in inducing on its own a weak anthocyanin synthesis under light, being its protein levels increased due to the release of its expression from the HY5 negative feedback regulation.

**Figure 9.**
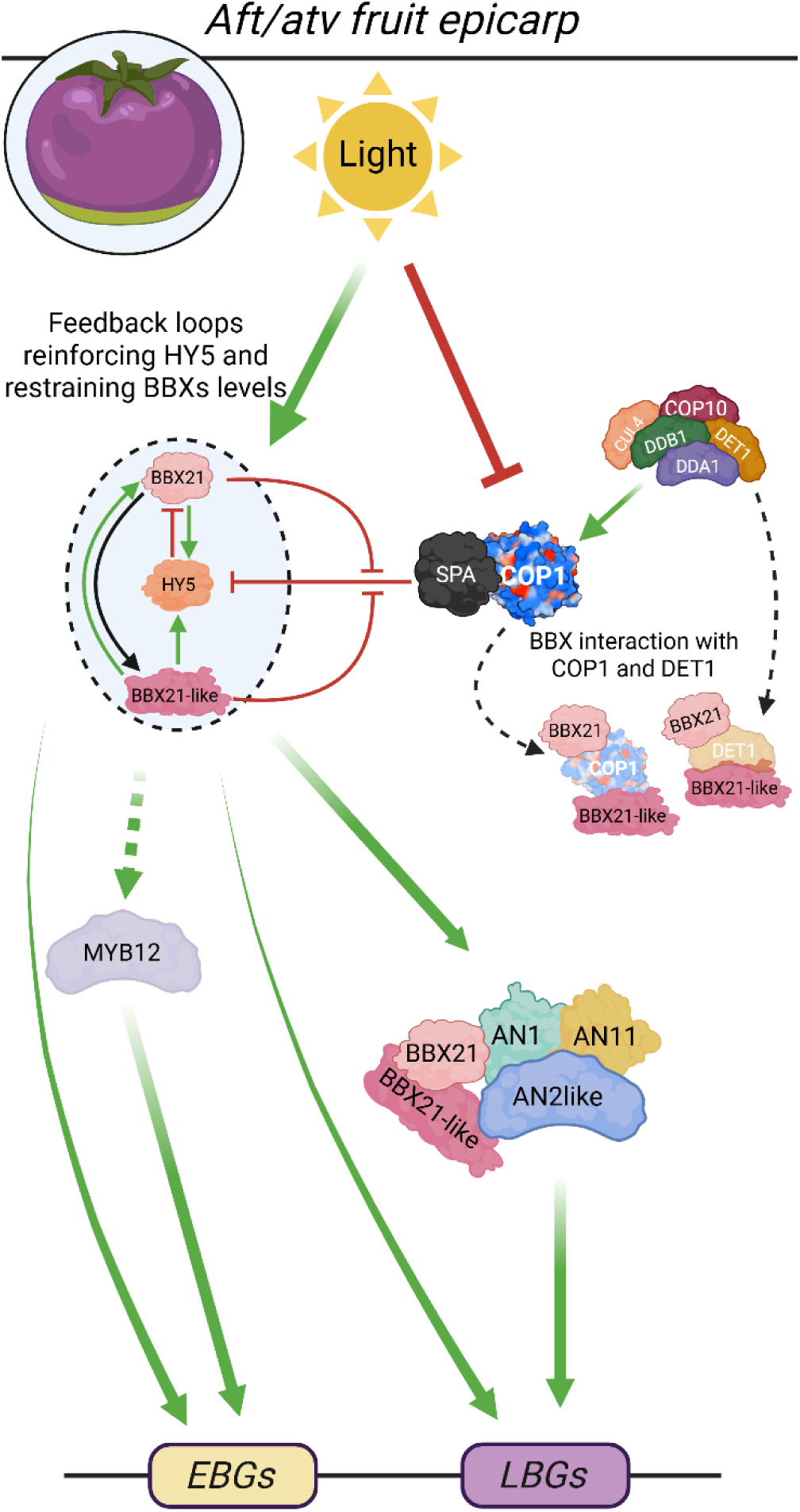
Recapitulating scheme of BBX-HY5 network in *Aft*/*atv* fruits.

**Figure 10.**
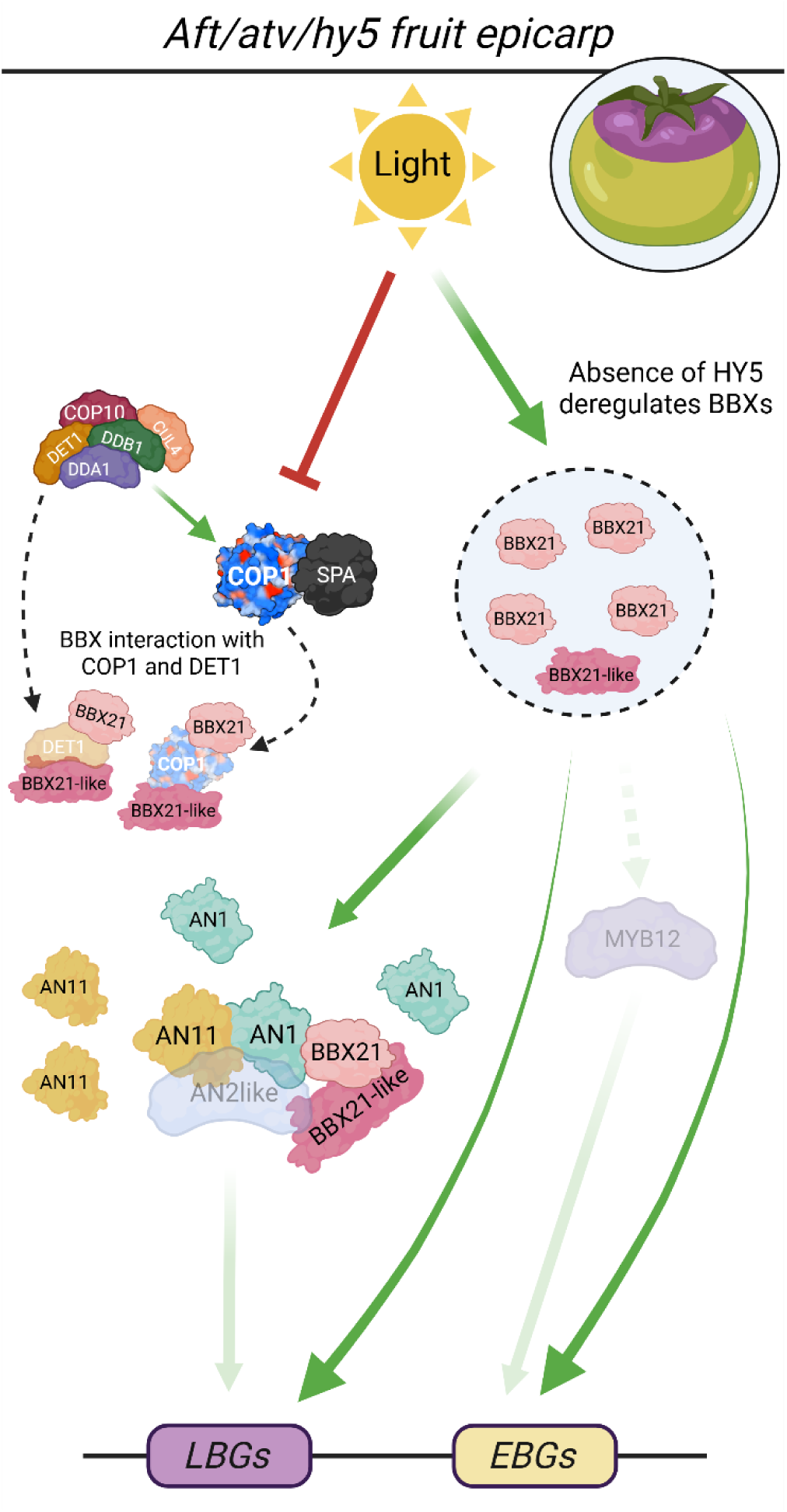
Recapitulating scheme of BBX network in *Aft*/*atv*/*hy5* fruits.

Altogether, the present results reveal novel light-responsive regulatory mechanisms of anthocyanin synthesis and accumulation in purple tomato fruits. The silencing of each of the two BBX proteins here studied may help in the future in unravelling their possible partial overlapping effects, finally disclosing the extent of functional redundance.

## Materials and Methods

### Plant materials and growth conditions

The tomato lines *Aft/Aft* x *atv/atv* (*Aft/atv*), *Aft/Aft* x *atv/atv x hy5/hy5* (*Aft/atv/hy5*) (Menconi et al. 2025) and *Aft/Aft* x *atv/atv* x *hp2/hp2* (*Aft/atv/hp2*) in the MicroTom background (Sestari et al. 2014) were used in the present research. Seeds were germinated in rock-wool plugs (Grodan, Canada). Two-week-old seedlings were transplanted in pots containing a 70:30 soil/expanded clay mixture and placed in a growth chamber with 23°/20°C day/night temperature, 12h photoperiod, 150 µmol photons m^-2^ s^−1^, and 40% relative humidity.

### Gene cloning and plasmid construction

Full length cds of tomato *Solyc12g089240* (*BBX21*), *Solyc04g081020* (*BBX21like*), and *Solyc01g056340* (*DET1*) were amplified by PCR using the “Phusion High-Fidelity DNA Polymerase” (Thermo Fisher Scientific, USA) and the oligonucleotide primers reported in Supplementary Table S1, starting from RNA extracted from the fruit peel using the “Spectrum Plant Total RNA Kit” (Merck, Germany), treated with DNase and reverse-transcribed with the SuperScript IV Reverse Transcriptase (Thermo Fisher Scientific). The full length cds of tomato *Solyc08g061130* (*HY5*), *Solyc12g005950* (*COP1 homolog*), *Solyc11g011980* (*COP1like X1 isoform*), *Solyc10g086250* (*AN2like*), *Solyc09g065100* (*AN1*) and the promoters of tomato *Solyc09g091510* (*CHS1*), *Solyc05g053550* (*CHS2*), *Solyc02g083860* (*F3H*), *Solyc02g085020* (*DFR*)*, Solyc10g086250* (*AN2like*), and *Solyc09g065100* (*AN1*) were produced in previous studies (Kiferle et al. 2015; Colanero et al. 2019; Menconi et al. 2025). The promoters of tomato *BBX21*, *BBX21like*, and *HY5* were obtained by gene synthesis (Thermo Fisher Scientific). The cds and the promoter sequences were individually cloned into pENTR/D-TOPO (Thermo Fisher Scientific) vectors and sequenced (Eurofins Genomics, Germany). The entry clones were recombined with different destination vectors, as described below, via Invitrogen™ Gateway™ recombination cloning technology (Thermo Fisher Scientific). Multiple sequence alignments were performed using ClustalW and MUSCLE (https://www.ebi.ac.uk/Tools/msa).

### Phylogenetic analysis

The analysis was performed on the Phylogeny.fr platform (http://www.phylogeny.fr/index.cgi) (Dereeper et al. 2008) using default programs and parameters. MAFFT was used for multiple alignment, and PhyML was used for phylogenetic analysis. Results were visualized using interactive Tree Of Life (iTOL) (https://itol.embl.de/) (Letunic and Bork, 2021)

### Protein structure modelling

Protein structures were assessed bioinformatically by using the SWISS-MODEL service (Bienert et al. 2017), operating through homology modelling. Briefly, aminoacidic sequences of the proteins of interested were used to automatically identify template models in the database available. Models of the proteins of interest were then built based on the most similar proteins found and the models with the highest accuracy were chosen.

### Anthocyanin quantification

50 mg of fruit peel were ground and extracted overnight in 300 µl of HCl 1% (v/v) in methanol. Extracts were recovered, diluted with 200 µl of distilled water, and one volume of chloroform was added to remove chlorophylls through mixing and centrifugation (2 minutes at 14,000 x g). Anthocyanin containing aqueous phase was recovered and, after addition of 600 µl of an aqueous solution containing 40% HCl 1% (v/v) in methanol, absorbances at 530 and 657 nm were determined spectrophotometrically and subtracted. Anthocyanins were expressed as mg petunidin-3-(p-coumaroyl)-rutinoside-5-glucoside g^-1^ fresh weight. Mean values were calculated from six biological replicates (obtained by pooling different fruits from different plants).

### Transient transformation of tomato protoplasts

Leaf protoplasts were isolated following the protocol in Shi et al. (2012) from 3-week-old MicroTom plants, grown as reported above. Polyethylene glycol-mediated protoplast transformation was carried out as in Yoo et al. (2007).

### Transactivation assays in tomato protoplasts

Transactivation assays by the dual-luciferase system were performed with the *Renilla reniformis* (Renilla) and *Photinus pyralis* (Firefly) luciferase (Luc) enzymes. The effector constructs *35S:BBX21* and *35S:BBX21like* were produced by recombining the *BBX21* and *BBX21like* entry clones in pENTR/D-TOPO vectors with the Gateway^TM^ compatible destination vector p2GW7 (Karimi et al. 2002). The reporter constructs *promoter_CHS1:Firefly_Luc, promoter_CHS2:Firefly_Luc, promoter_F3H:Firefly_Luc, promoter_AN2like:Firefly_Luc, promoter_AN1:Firefly_Luc, promoter_BBX21:Firefly_Luc, promoter_BBX21like:Firefly_Luc,* and *promoter_HY5:Firefly_Luc* were produced by recombining the sequences of the promoters cloned and ligated in pENTR/D-TOPO vectors, as above described, with the Gateway^TM^ compatible destination vector pPGWL7 (Karimi et al. 2002) containing the Firefly *Luc* gene. The effector constructs *35S:AN2like* and *35S:AN1* and the reporter construct *promoter_DFR:Firefly_Luc* were produced in previous studies (Kiferle et al. 2015; Colanero et al. 2019). A *35S:Renilla_Luc* vector (Kiferle et al. 2015) was used to normalize luminescence values detected in protoplasts. Effector and reporter plasmids were co-transfected in protoplasts (5 µg for each effector plasmid, 5 µg for the reporter construct and 1 µg for the *35S:Renilla_Luc*). After an overnight incubation under dark, protoplasts were lysed and Firefly and Renilla luciferase activities were sequentially measured through the “Dual-Luciferase® Reporter Assay System” (Promega, USA) with a Lumat LB 9507 Tube Luminometer (Berthold Technologies, USA). Data were expressed as relative luciferase units (RLU = Firefly_Luc/Renilla_Luc).

### Yeast One Hybrid assay

Yeast One Hybrid assays were performed with the “Grow ‘n’ Glow One-Hybrid System” (Mobitech, Germany) according to the manufacturer instructions, with the modifications described in a previous study (Menconi et al. 2025). Briefly, the prey constructs *AD-BBX21* and *AD-BBX21like* were produced by recombining the entry vector containing their cds into the destination vector pJG4-5_GW. The reporter constructs *pBBX21:GFP, pBBX21like:GFP,* and *pHY5:GFP* were produced by recombining the entry vector containing the relative promoter sequences into the destination vector pGNG2_GW. The prey construct *AD-HY5* and the reporter constructs *pCHS1:GFP, pCHS2:GFP, pF3H:GFP, pDFR:GFP, pAN2like:GFP* and *pAN1:GFP* were produced in a previous study (Menconi et al. 2025). Different combinations of prey and reporter constructs were transformed into *Saccharomyces cerevisiae* strain EGY48 and successful transformants screened on -ura -trp synthetic medium containing galactose and raffinose. Plates were visually inspected under UV light for GFP signal. For quantitative Y1H assay each transformed yeast strain obtained was inoculated in liquid media and grown overnight at 30°C, then back diluted to OD 1.0, centrifuged at 15 000 x g for 30 s, the supernatant discarded, and the pellet resuspended in sterile water to a final OD of 1.0. The GFP level of each yeast strain was quantified using a TriStar Microplate Reader (Berthold Technologies).

### RNA isolation, cDNA synthesis, and quantitative RT-PCR analysis

Total RNA extracted from tomato fruit peel with the “Spectrum^TM^ Plant Total RNA Kit” (Merck) was subjected to DNase treatment and then reverse transcribed into cDNA using the “Maxima First Strand cDNA Synthesis Kit for RT-qPCR, with dsDNase” (Thermo Fisher Scientific). Quantitative RT-PCR was performed with a QuantStudio 3 Real-Time PCR system (Thermo Fisher Scientific) using the “PowerUp™ SYBR® Green Master Mix” (Thermo Fisher Scientific) and the primers listed in Supplementary Table S2. Tomato *Elongation Factor 1-alpha* (*EF1A*) and *ACTIN*-2 (*ACT*-2) were used as reference genes. The relative quantification of gene expression was performed with the double delta Ct (ΔΔCt) method using the geometric averaging of the reference genes.

### Cellular localization of the candidate factors

The *BBX21*, *BBX21like*, and *HY5* entry vectors were recombined with the Gateway^TM^ compatible destination vector p2FGW7 (Karimi et al. 2002). Protoplasts were isolated as described, transformed with 5 µg DNA for each plasmid, and incubated in the dark at 23°C for 16h before subsequent analysis. The *35S:NLS-mCherry* construct was used as nuclear marker (Menconi et al. 2025). Fluorescence for GFP and for RFP were imaged with a Nikon Eclipse Ti-5 video-confocal microscope (Nikon Corporation, Japan) using suitable filters.

### Tomato stable transformation and regeneration

The overexpression vectors were generated by recombining the entry clones containing the cds of *BBX21* and *BBX21like* with the Gateway™ compatible binary vector pK7WG2 (Karimi et al. 2002). *Agrobacterium tumefaciens* cells from the GV3101 (MP90) strain hosting the different constructs were incubated with tomato cotyledons and young leaves of MicroTom *Aft*/*atv* plants, which were transformed following the protocol described in Zuluaga et al. (2008). Transgenic regenerating plants were selected for their kanamycin resistance and the presence of the pK7WG2 construct (Supplementary Table S3). For RNA extraction and qPCR analyses, peel from fruits at the MG stage isolated from different independent transgenic lines were used.

### Yeast Two Hybrid assay

Yeast Two Hybrid assays were performed with the ProQuest Two-Hybrid System (Thermo Fisher Scientific) according to the manufacturer instructions. Bait constructs pDEST32_*BBX21* and pDEST32_*BBX21like* were produced by recombining the entry vector containing their cds into the destination vector pDEST32_GW. The prey constructs pDEST22_*BBX21*, pDEST22_*BBX21like*, pDEST22_*HY5*, pDEST22_*AN2like*, pDEST22_*AN1*, pDEST22_*DET1*, pDEST22_*COP1like X1 isoform*, pDEST22_*COP1 homolog* were produced by recombining the entry vector containing the relative cds into the destination vector pDEST32 GW. The bait/prey combinations pEXP32_Krev1/pEXP22_RalGDS-wt and pEXP32_Krev1/pEXP22_RalGDS-M2 were used as positive and negative controls, according to the manufacturer instructions. Different combinations of prey and reporter constructs were transformed into *S. cerevisiae* strain MAV203 and successful transformants screened on -leu -his -trp synthetic medium containing 25 mM 3-amino-1,2,4-triazole (3-AT; Merck) and on -leu -his synthetic medium containing X-alpha-Gal (Takara Bio, Japan).

### Protein extraction and Western blot immunostaining

Protein extraction and cellular fractionation were performed as described by Law et al. (2013) with adjustments by Menconi et al. (2025), starting from 1 g of frozen plant material of fruits or leaves. Total protein content was quantified using the ‘Pierce BCA Protein Assay Kit’ (Thermo Fisher Scientific). A total of 25 micrograms of protein extracts were denatured at 95°C for 5 min in XT Buffer (Bio-Rad, USA) supplemented with 0.8 M DTT. Denatured extracts were loaded onto polyacrylamide gels (Thermo Fisher Scientific), separated by SDS-PAGE, and semi-wet-transferred onto a PVDF membrane (Bio-Rad) with the TransBlot Turbo Transfer System (Bio-Rad). The membranes were then incubated with rabbit polyclonal anti-HY5 (PHY0748S -PhytoAB, USA) or anti-H3 (E-AB-22081 -Elabscience, USA) antibodies, all diluted 1:1000. HRP-conjugated goat anti-rabbit IgG (AS09 602 -Agrisera, Sweden) or HRP-conjugated rabbit anti-mouse IgG (AP160A, Merck) were used as the secondary antibodies. Clarity Max Western ECL Substrate (Bio-Rad) and a ChemiDoc MP Imaging System (Bio-Rad) were used to detect signals. Image Lab software (Bio-Rad) was used to quantify the intensity of the detected bands. The HY5 quantification data from each biological replicate were normalized to the bands corresponding to histone H3, which was used as a housekeeping.

### Statistics

Statistical analyses were performed with GraphPad Prism 10.00 (www.graphpad.com/scientific-software/prism/). Data were analysed by t-test or one-way ANOVA with differences measured using the Tukey’s honest significant difference (HSD) multiple comparisons test.

### Figure preparation

Figures were created with BioRender.com

## Acknowledgments

We thank Dr. Giacomo Novi for his expertise and assistance with plant growth and Dr. Francesco Fioriti for his assistance in microscopic imaging. We also thank Prof. L.E.P. Peres (Universidade de São Paulo, Brazil) for providing seeds of the “triple mutant” line *Aft*/*atv*/*hp2* and Sebastien Santini (CNRS/AMU IGS UMR7256) and the PACA Bioinfo platform for the availability and management of the phylogeny.fr website used.

## Author contributions

J.M. and S.G. conceived the project. S.G. and P.P. supervised the project. J.M. performed most of experiments and analyses. N.L.M. carried out the yeast one hybrid and two hybrid assays. J.M. wrote the manuscript and prepared the figures. S.G. and P.P. revised the manuscript. All authors reviewed and approved the final manuscript.

## Supplementary data

The following materials are available in the online version of this article:

**Supplementary Table S1.** List of oligonucleotide primers used for CDS cloning.

**Supplementary Table S2.** List of oligonucleotide primers used for qPCR analysis.

**Supplementary Table S3.** List of oligonucleotide primers used for genotyping tomato transgenic lines overexpressing *BBX21* or *BBX21like*.

**Supplementary Figure S1.** Anthocyanin biosynthetic pathway and its regulation.

**Supplementary Figure S2.** Phylogenetic analysis of BBX gene family in *Solanum lycopersicum* and *Arabidopsis thaliana*.

**Supplementary Figure S3.** Promoter sequences of anthocyanin and light-related genes analysed in the Yeast One Hybrid (Y1H) assay.

**Supplementary Figure S4.** Yeast One Hybrid assay.

**Supplementary Figure S5.** Phenotypes of the young MicroTom plants of all the transgenic lines overexpressing the selected BBX factors.

**Supplementary Figure S6.** Phenotypes of representative young MicroTom plants overexpressing the selected BBX factors.

**Supplementary Figure S7.** *BBX21* or *BBX21like* overexpression affects anthocyanin biosynthetic genes in tomato fruits.

**Supplementary Figure S8.** *BBX21* or *BBX21like* overexpression affects anthocyanin regulatory and light-related genes in tomato fruits.

**Supplementary Figure S9.** Negative controls for Yeast Two Hybrid assay.

**Supplementary Figure S10.** Representative Western blot showing HY5 protein levels in fruits from *Aft*/*atv* lines overexpressing the two BBX factors.

**Supplementary Figure S11.** Representative Western blot of HY5 protein levels in leaves from *Aft*/*atv* lines overexpressing the two BBX factors.

**Supplementary Figure S12.** Phenotypes of young *Aft/atv/hy5* plants compared with background plants.

**Supplementary Figure S13.** Knockout of HY5 affects anthocyanin and light-related genes in tomato fruits at different growth stages.

## Data availability

The data that supports the findings of this study are available in the supporting material of this article.

